# Age-Related Remodeling of Cross-Tissue Transcriptional Coordination in the Human Motor System

**DOI:** 10.64898/2026.09.21.753254

**Authors:** Huan Nie, Kunihiro Sakuma

## Abstract

How aging is coordinated across the anatomically distinct tissues that collectively constitute the human motor system remains unclear. Here, we integrated data from the Genotype-Tissue Expression (GTEx) project and Gene Expression Omnibus (GEO), comprising 7,145 samples from 15 human tissues spanning three functional levels of motor control, signal transmission, and peripheral execution, and reconstructed age-related transcriptional trajectories using generalized additive models. Most age-related genes were shared across multiple tissues, yet their direction of regulation, effect magnitude, and temporal trajectories showed marked tissue specificity. Despite the clear central-to-peripheral functional organization of the motor system, transcriptional aging did not follow a stable temporal sequence or spatial gradient along this functional chain. Instead, cross-tissue transcriptional synchrony progressively increased from early to mid-adulthood, reached a relatively high level around midlife, and subsequently declined in later life, a pattern supported at both the gene and pathway levels. This late-life loss of coordination was selective, prominently involving the peripheral nerve-skeletal muscle axis and multiple tissue connections involving the striatum. Our analyses indicate that transcriptional aging of the human motor system is characterized by tissue-specific responses built upon a broadly shared molecular basis of aging, together with age-dependent reorganization of cross-tissue coordination and selective late-life decoupling.

## Introduction

Motor function depends on the coordinated activity of anatomically distinct but functionally interconnected components of the nervous and musculoskeletal systems, including cortical and basal ganglia circuits involved in motor control, brainstem and spinal pathways involved in signal transmission, and peripheral nerves and skeletal muscle responsible for motor execution [1,2]. During aging, muscle strength and physical performance decline, accompanied by age-related changes in sensorimotor control and the neural circuits supporting movement [3,4]. Understanding whether different motor-related tissues age independently or remain molecularly coordinated over the life course, and whether this coordination changes with age, is therefore important for elucidating motor-system aging.

Aging induces widespread transcriptional changes, but the extent and magnitude of these changes differ markedly across tissues [5]. Multi-organ studies have further revealed organ-specific temporal patterns of molecular aging, indicating that different tissues do not necessarily undergo age-related remodeling at the same rate or at the same stages of life [6]. At the same time, subsets of age-related genes and biological processes are shared across multiple tissues, and synchronized age-associated gene-expression changes have been observed across human tissues [6,7]. More broadly, altered intercellular communication is recognized as a hallmark of aging [8], while studies of human organ systems have demonstrated substantial heterogeneity as well as coordinated or interrelated patterns of aging across organs [9,10]. Together, these observations suggest that tissue aging is neither completely independent nor uniformly synchronized. An important question is therefore whether age-related molecular changes remain coordinated across functionally connected tissues and whether this coordination itself changes over the life course.

This question is particularly relevant to the motor system. Motor control, neural transmission, and muscular execution form a functionally interconnected system [1,2], but it remains unclear whether this functional organization corresponds to a temporal organization of molecular aging. Specifically, it is unknown whether central, transmission, and peripheral tissues undergo major transcriptional changes in a sequential manner along the motor functional chain, or whether functionally adjacent tissues exhibit more similar age-related responses. Previous studies have shown that transcriptional coordination can change with age, including changes in gene– gene relationships within tissues [11,13] and coordinated expression relationships across tissues [7,12]. However, whether such coordination is dynamically reorganized across the anatomically and functionally connected tissues of the human motor system remains unclear. In particular, it is unknown whether cross-tissue transcriptional coordination follows a characteristic trajectory across adulthood, whether it selectively weakens in later life, and which tissue connections and biological processes are most affected.

Here, we integrated data from the Genotype-Tissue Expression (GTEx) project [14] with multiple independent human transcriptomic datasets from the Gene Expression Omnibus (GEO) [15] to reconstruct age-related transcriptional trajectories across 15 tissues spanning three functional levels of the motor system: motor control, signal transmission, and peripheral execution. We first compared the extent, direction, trajectory pattern, and sharing of age-related transcriptional changes across tissues and tested whether these changes followed a stable temporal sequence or spatial organization along the motor functional chain. We then quantified the synchrony of gene-expression changes and pathway responses across tissues within consecutive age windows to determine how cross-tissue coordination is reorganized with age and to identify tissue connections and functional processes showing pronounced divergence. Through this study, we investigated transcriptional aging of the human motor system from the perspective of age-dependent changes in cross-tissue coordination rather than isolated changes within individual tissues.

## Methods

### Data sources and processing

Raw read counts from the GTEx project V11 were used. Sample metadata were matched by sample ID, including age, sex, RNA integrity number (SMRIN), ischemic time (SMTSISCH), and Hardy scale of death (DTHHRDY). The midpoint of each 10-year interval was used as the age value. For each tissue, lowly expressed genes were filtered using filterByExpr in edgeR [16], followed by trimmed mean of M-values (TMM) normalization [17] and transformation to log counts per million (logCPM). Gene expression values were then Z-score standardized within each study.

GEO datasets included human microarray and bulk RNA-seq studies with individual-level age information. Each GSE dataset was processed independently. For microarray datasets, probe identifiers were mapped to gene symbols using the corresponding platform annotation, and expression values were averaged when multiple probes mapped to the same gene. For RNA-seq datasets, gene identifiers were converted to gene symbols, lowly expressed genes were filtered, and expression data were TMM-normalized and transformed to logCPM. All GSE datasets were subsequently Z-score standardized by gene within each study.

Gene symbols and tissue names were harmonized across GTEx and GEO. Each GTEx tissue and each individual GSE dataset were treated as a separate study. Within each tissue, the union of genes detected across studies was retained. For genes not measured in all studies, model fitting used only samples with observed expression values. To harmonize the age scale between GTEx and GEO, exact ages in GEO were grouped into 10-year intervals corresponding to the GTEx age bins, and the midpoint of each interval was used as the age value.

### Age-related transcriptional trajectories

Genes were analyzed only if they had at least 10 valid samples, an age span of at least 30 years, and coverage of at least three age groups. For each gene within each tissue, an age-related expression trajectory was fitted using a Gaussian generalized additive model (GAM) [18,19]. Standardized gene expression was used as the response variable, and age was modeled using a thin-plate regression spline, with the basis dimension set to a maximum of 5. Sex was included as a fixed effect and study as a random intercept. When a study contained multiple platforms and/or subtissues, platform and/or subtissue were additionally included as random intercepts nested within study. GTEx samples were further adjusted for SMRIN, SMTSISCH, and DTHHRDY. Models were estimated using restricted maximum likelihood (REML).

The age effect was assessed using the GAM smooth term for age, and *P*-values were corrected within each tissue using the Benjamini–Hochberg procedure. Genes with a false discovery rate (FDR) < 0.05 were defined as age-related genes (ARGs). The effective degrees of freedom (edf) of the age smooth were used to describe trajectory complexity, with edf ≥ 1.5 classified as a nonlinear trajectory.

To characterize the overall direction of an age-related transcriptional trajectory, the net age-related change was defined as the difference between the predicted GAM age effects at the oldest and youngest available age groups:

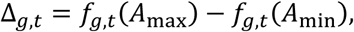

where *g* denotes gene, *t*denotes tissue, *f_g_*_,*t*_ is the GAM-estimated smooth age effect, and *A*_min_ and *A*_max_ are the youngest and oldest available age groups for that gene in that tissue. A positive value (Δ > 0) indicated an overall increase with age, whereas a negative value (Δ < 0) indicated an overall decrease.

### Cross-tissue similarity of age effects

To compare overall transcriptional aging patterns across tissues, a directional age-effect ranking was calculated for each successfully fitted gene:

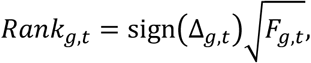

where *F_g_*_,*t*_ is the *F* statistic of the GAM smooth age term.

For each pair of tissues, Spearman correlation coefficients (*ρ*) were calculated between their rank vectors using genes that were successfully modeled in both tissues. These correlations were used to quantify the similarity of overall age-related transcriptional effects between tissues. For ARGs that reached FDR < 0.05 in both tissues, directional concordance was defined as:

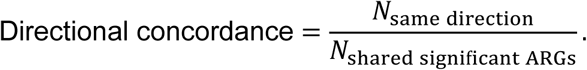

### Age windows of transcriptional change and cross-tissue synchrony

The fitted GAM age smooths were used to predict gene expression effects at consecutive 10-year age points. The transcriptional change within each age window was defined as the difference between predictions at two adjacent age points:

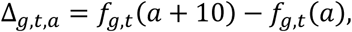

where *a* is the starting age of the corresponding 10-year window.

System-level analyses were restricted to the age range shared by all tissues and to genes with valid estimates in every tissue and every age window. Within each age window, Spearman correlation coefficients were calculated between the gene-level Δvectors of each tissue pair. These correlations were used as measures of cross-tissue transcriptional synchrony. Mean *ρ*values were then summarized across all tissue pairs, for individual tissues, and within and between functional levels to characterize age-related changes in cross-tissue synchrony. A gene-level bootstrap was used to assess the robustness of synchrony estimates to gene sampling. Common genes were sampled with replacement for 1,000 iterations. Tissue-pair correlations and system-wide mean synchrony were recalculated in each iteration, and the 2.5th and 97.5th percentiles of the bootstrap distribution were used to define 95% intervals. For analyses of functional-level ordering and the relationship between functional distance and transcriptional age effects, tissue labels were permuted to generate null distributions, and two-sided permutation *P*-values were calculated. For each tissue pair, the difference between late-life and early-to-mid-adulthood synchrony was calculated as:

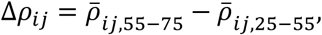

where *ρ̅ _ij_*_,55−75_ and *ρ̅ _ij_*_,25−55_ represent the mean synchrony between tissues *i*and *j*across the 55–75-year and 25–55-year age windows, respectively.

### Pathway enrichment and cross-tissue functional synchrony

Pre-ranked gene set enrichment analysis was performed using the Human Molecular Signatures Database (MSigDB) [20] C5 Gene Ontology Biological Process (GO-BP) gene sets. Normalized enrichment scores (NESs) and multiple-testing-adjusted FDR values were calculated using fgseaMultilevel. GO-BP terms with FDR < 0.05 were considered significantly enriched. For analyses across the full age range, genes were ranked using the directional GAM age-effect statistic described above, and gene sets containing 10∼1,000 genes were tested. For age-window analyses, genes were ranked according to their corresponding window-specific Δvalues, and gene sets containing 10∼500 genes were tested. Cross-tissue functional synchrony within each age window was quantified using the same framework as the gene-level analysis, with pathway NES values used as pathway-level effect sizes. Spearman correlations and directional concordance were calculated between tissues to quantify the similarity of pathway responses.

### Semantic modules

A GO-BP term was defined as a robust layer-level process when it met three criteria: it was significant at FDR < 0.05 in at least 50% of the available tissues within that functional level, it was supported by at least two tissues, and at least 75% of the significant tissues showed the same NES direction. Functional-level analyses were restricted to the age range shared by all tissues. Pairwise semantic similarity between robust GO-BP terms was calculated using the Wang method. An undirected edge was placed between two GO terms when their Wang semantic similarity exceeded 0.7, generating a GO semantic-similarity network [21]. Connected components were first identified. Components containing at least three nodes and at least one edge were further partitioned using edge-betweenness community detection [22], whereas components containing two or fewer nodes, or no edges, were retained as individual modules. For each module, an overall direction was assigned according to the directions of its member GO-BP terms. Modules in which at least 75% of member GO-BP terms agreed in direction were defined as coherent functional modules.

## Results

### Data Sources

We included 7,145 tissue samples from 49 independent studies spanning 15 tissues including frontal/prefrontal cortex (PFC), cortex (CTX), somatosensory cortex (SSC), putamen (PUT), caudate (CAU), nucleus accumbens (NAc), substantia nigra (SN), thalamus (THA), cerebellum (CBL), pons (PON), medulla (MED), central white matter (CWM), spinal cord (SC), tibial nerve (TN), and skeletal muscle (SKM).The samples covered ages 18∼106 years, with tissue-specific sample sizes ranging from 53 to 1,587 and the number of genes available for analysis ranging from 17,184 to 48,849 (Fig. 1A–C).

**Fig. 1:**
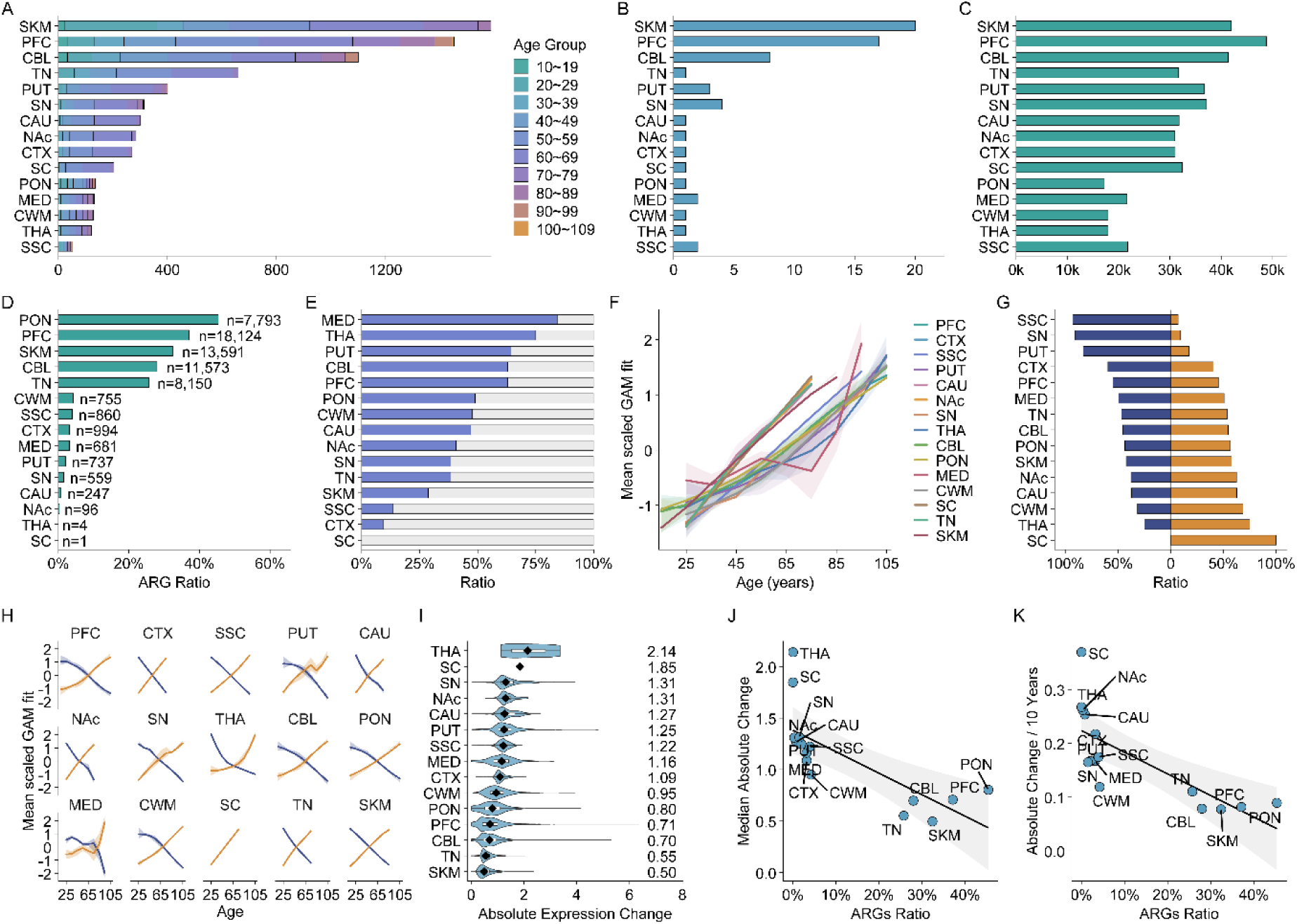
Multidimensional heterogeneity of age-related transcriptional changes across tissues of the human motor system. **A** Sample distribution across 10-year age groups in the 15 motor-system tissues. **B** Number of independent studies included for each tissue. **C** Number of genes tested by GAM in each tissue. **D** Proportion of age-related genes (ARGs) among all tested genes; numbers indicate ARG counts. **E** Proportions of ARGs with linear (edf < 1.5) and nonlinear (edf ≥ 1.5) age trajectories. **F** Direction-aligned GAM trajectories of ARGs across tissues; lines indicate mean scaled GAM fits and shaded regions indicate interquartile ranges. **G** Proportions of ARGs showing overall positive (Up) or negative (Down) age-related changes. **H** GAM trajectories of Up and Down ARGs across tissues; lines indicate mean scaled GAM fits and shaded regions indicate interquartile ranges. **I** Distribution of absolute age-related expression changes (|Δ|) among ARGs; black diamonds indicate tissue medians. **J** Relationship between ARG proportion and median absolute age-related expression change. **K** Relationship between ARG proportion and median absolute expression change normalized per 10 years of age span. In **J** and **K**, solid lines indicate linear regression fits and shaded regions indicate 95% confidence intervals.

### Multidimensional heterogeneity of age-related transcriptional changes

GAM analysis revealed marked differences in the extent of age-related transcriptional changes across tissues. PON, PFC, SKM, CBL, and TN showed both higher numbers and proportions of age-related genes (ARGs; FDR < 0.05) than most other tissues (Fig. 1D). When the analysis was restricted to the 11,208 genes detected across all 15 tissues, PFC, CBL, SKM, PON, and TN retained relatively high numbers and proportions of ARGs (Supplementary Fig. 1A). By contrast, very few ARGs were detected in SC and THA (Fig. 1D). Age-related trajectory shapes also differed substantially across tissues in their degree of linearity and nonlinearity (Fig. 1E, F, H; Supplementary Fig. 1B). For example, 84.3% of ARGs in MED showed nonlinear trajectories, whereas only 9.3% of ARGs in CTX, which had a similar overall extent of age-related transcriptional change, were nonlinear (Fig. 1E; Supplementary Fig. 1B). Pronounced differences were also observed among tissues with relatively broad age-related effects. In SKM, 71.3% of ARGs followed approximately linear trajectories, whereas 62.9% of ARGs in both PFC and CBL were nonlinear (Fig. 1E; Supplementary Fig. 1B).

The direction of age-related regulation was also strongly tissue specific, with no uniform transcriptional direction across the motor system (Fig. 1G, H). SSC, SN, and PUT were biased toward age-related decreases, whereas CWM, CAU, and NAc were biased toward increases. MED, TN, and CBL showed more balanced proportions of increasing and decreasing ARGs (Fig. 1G, H). The breadth and magnitude of age-related effects were also markedly dissociated. Although SKM and TN showed relatively high proportions of ARGs, their ARGs exhibited the smallest expression changes. Conversely, PUT, CAU, NAc, and SN showed lower proportions of ARGs but larger effect magnitudes. PUT and SN were particularly notable for combining relatively large expression changes with a pronounced bias toward age-related decreases (Fig. 1D, G, I).

Tissue sample size was moderately positively correlated with the number of ARGs (Spearman ρ = 0.543, *P* = 0.037; Supplementary Fig. 1C), but was not significantly associated with either the proportion of ARGs (ρ = 0.368, *P* = 0.177; Supplementary Fig. 1D) or the proportion of nonlinear ARGs (ρ = −0.039, *P* = 0.889; Supplementary Fig. 1E). Both the proportion of ARGs and the proportion of nonlinear ARGs were positively associated with the span of age coverage (ρ = 0.516, *P* = 0.049 and ρ = 0.614, *P* = 0.015, respectively; Supplementary Fig. 1F, G). The proportion of ARGs was strongly inversely correlated with the median absolute expression change among significant ARGs (ρ = −0.914, *P* < 0.001), and this relationship remained after normalization for age span (ρ = −0.904, *P* < 0.001; Fig. 1J, K). Together, these results indicate that sample size may partly influence the number of detected ARGs but does not fully account for differences in ARG proportions or trajectory nonlinearity, and that different tissues may exhibit distinct transcriptional aging patterns characterized by either broad, low-amplitude changes or more restricted, high-amplitude changes.

### Broad cross-tissue sharing of ARGs

In contrast to the marked tissue-level heterogeneity, ARGs showed extensive cross-tissue sharing when analyzed against the common gene background (Fig. 2A). Overall, 87.1% of ARGs were significant in at least two tissues, with some shared across as many as nine tissues, whereas only 12.9% were significant in a single tissue (Fig. 2B). Within individual tissues, the vast majority of ARGs were shared with at least one other tissue, and tissue-specific ARGs accounted for no more than 5% of ARGs in any tissue (Fig. 2C). Thus, although tissues exhibited distinct transcriptional aging profiles, these differences did not arise from entirely independent sets of ARGs but rather from a broadly shared age-related transcriptional basis.

**Fig. 2:**
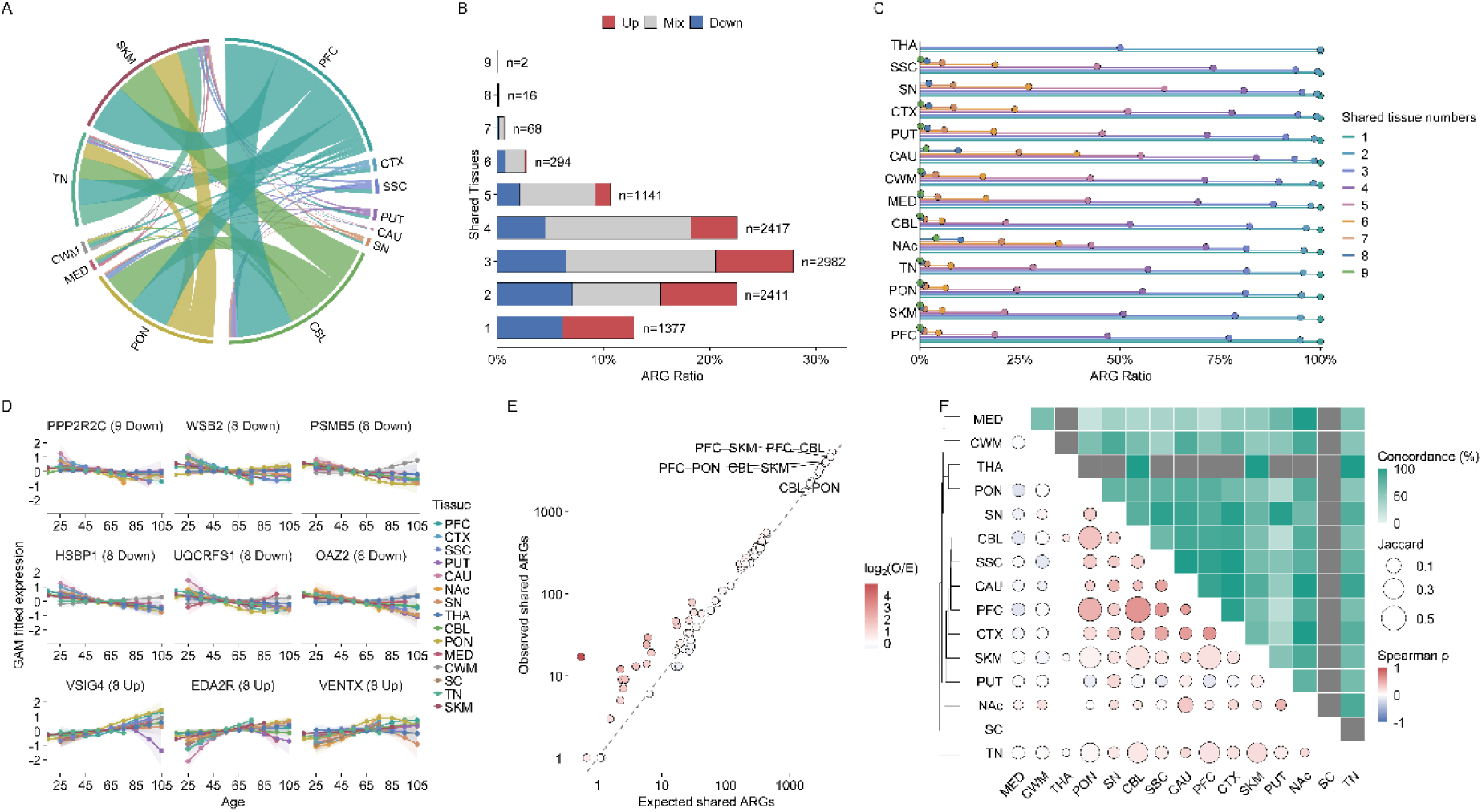
Broad cross-tissue sharing of age-related genes and tissue-specific age responses in the motor system. **A** Cross-tissue sharing of ARGs among the 15 tissues using the common-gene background. Only tissue pairs sharing ≥50 ARGs are shown. **B** Distribution of ARG sharing breadth and direction of age-related change; numbers indicate the number of ARGs shared across the corresponding number of tissues. **C** Cumulative proportions of ARGs in each tissue according to the number of tissues in which they were shared. **D** Age trajectories of representative core ARGs showing highly consistent directions across tissues. **E** Comparison of observed and randomly expected numbers of shared ARGs for each tissue pair. O and E denote the observed and expected numbers of shared ARGs, respectively. **F** Integrated pairwise matrix of ARG sharing and similarity of age-related responses. The upper triangle shows directional concordance among ARGs significant in both tissues, whereas the lower triangle shows Jaccard similarity of ARG sets and Spearman correlation coefficients (ρ) of overall age effects.

A small subset of core ARGs showed particularly consistent directions of change across tissues. PPP2R2C [23] decreased with age across nine tissues spanning central to peripheral components of the motor system, whereas EDA2R [24] consistently increased across eight tissues. UQCRFS1 [25] also showed a stable cross-tissue pattern; it encodes a subunit of mitochondrial complex III involved in mitochondrial respiration (Fig. 2D). These directionally conserved ARGs represent candidate markers of shared transcriptional aging across the motor system.

Despite this extensive sharing, the manner in which tissues responded to age differed substantially. PFC–CBL and PFC–SKM showed the highest ARG overlap (Jaccard = 0.565 and 0.485, respectively), and both overlaps exceeded random expectation (O/E = 1.13 and 1.06; Fig. 2E, F). Among shared ARGs, PFC–CBL showed 86.4% directional concordance and relatively high similarity in overall age effects (ρ = 0.607). By contrast, although PFC–SKM shared a large number of ARGs, their directional concordance was lower at 61.3%, and their overall age-effect similarity was only ρ = 0.169 (Fig. 2F). Thus, similar ARG composition across tissues could correspond to markedly different age-related response patterns.

This divergence was also evident among anatomically or functionally related tissues. PFC showed relatively high overall age-effect similarity with CTX, SSC, and CBL (ρ > 0.6), whereas CWM showed correlations close to zero with most tissues, and MED showed negative correlations with several tissues. Similarity was also relatively low for SC–TN across the transmission–peripheral interface and for TN–SKM within the peripheral level (ρ < 0.25; Fig. 2F). Together, these findings indicate that broadly shared ARGs provide a common molecular basis of aging, while individual tissues retain distinct responses to these shared age-related signals.

### Temporal ordering of transcriptional aging

Because aging progresses over time whereas the motor system is functionally organized from central control to peripheral execution [1,2], we further examined whether these two forms of progression corresponded to each other. First, we grouped all tissues into three functional levels: Control (PFC, CTX, SSC, PUT, CAU, NAc, SN, THA, and CBL), Transmission (PON, MED, CWM, and SC), and Peripheral (TN and SKM). Using the common-gene background across tissues, we tested whether transcriptional aging followed a temporal sequence across these functional levels. If aging progressed sequentially from central to peripheral tissues, the timing of major transcriptional changes would be expected to show a corresponding monotonic pattern. Based on the age at which cumulative transcriptional change reached 50% (T50), the mean T50 values were 50.6, 50.5, and 49.2 years for the Control, Transmission, and Peripheral levels, respectively (Fig. 3A, B). Functional level was not significantly associated with T50 (Spearman ρ = −0.073, permutation *P* = 0.798; Fig. 3B), providing no evidence for a stable temporal sequence from central to peripheral tissues or in the reverse direction.

**Fig. 3:**
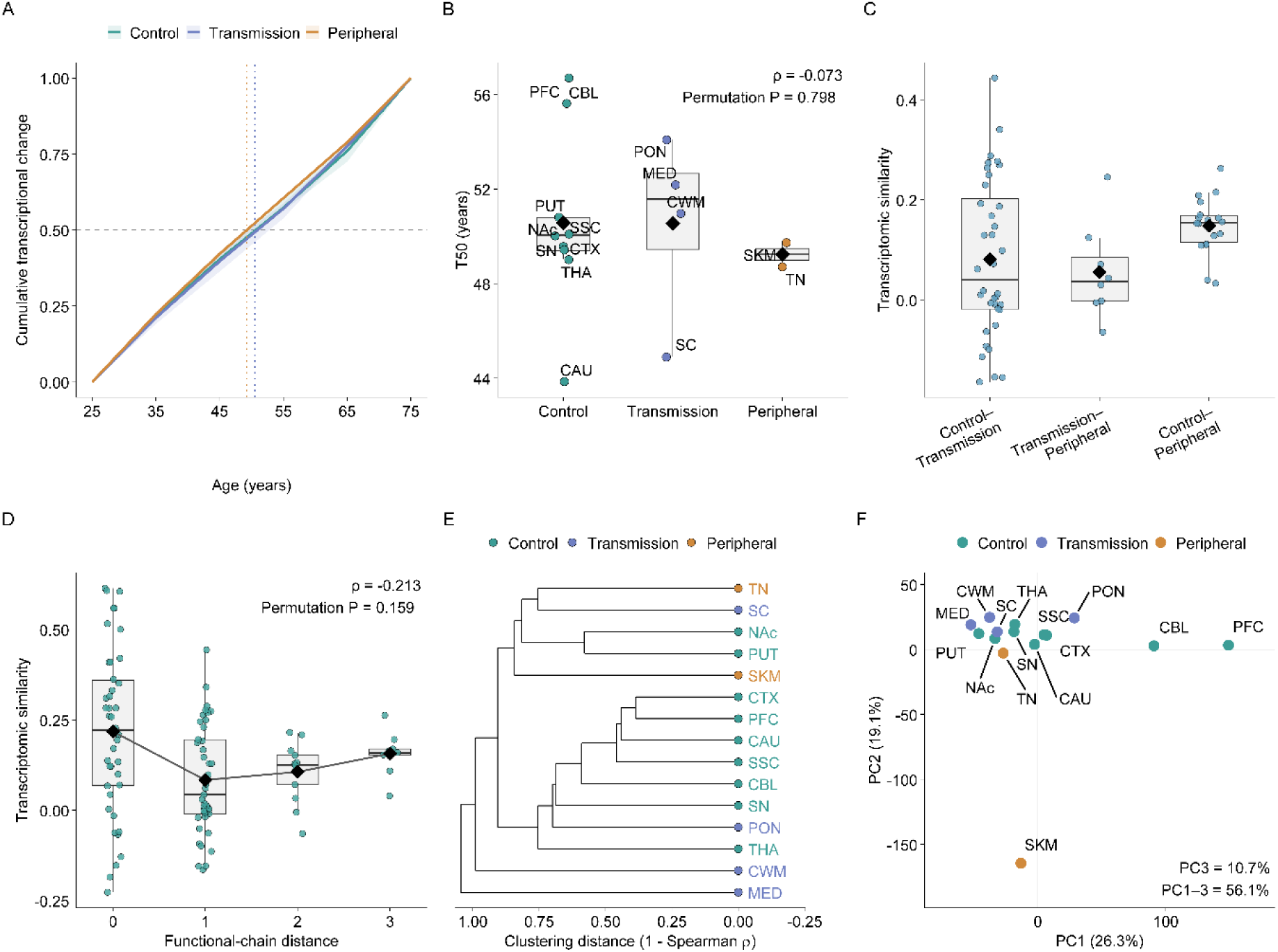
Temporal ordering of transcriptional aging and organization along the motor functional chain. **A** Cumulative transcriptional change across age for the three functional levels. **B** Distribution of tissue-specific T50 values across the Control, Transmission, and Peripheral levels. **C** Overall age-effect similarity between pairs of functional levels. **D** Relationship between functional-chain distance and cross-tissue age-effect similarity; distances of 0–3 represent progressively greater separation along the functional chain. **E** Hierarchical clustering based on overall age-effect similarity (1 − Spearman ρ). **F** Principal component analysis of overall age effects.

We next examined whether transcriptional aging exhibited a continuous organization consistent with the motor functional chain. If such a structure existed, age effects would be expected to be more similar between adjacent functional levels and least similar between more distant levels. However, the mean similarities for Control–Transmission (ρ = 0.081), Transmission–Peripheral (ρ = 0.055), and Control–Peripheral (ρ = 0.148) did not show the expected gradient (Fig. 3C). Functional-chain distance showed only a weak negative correlation with age-effect similarity (ρ = −0.213), which was not significant by permutation testing (*P* = 0.159; Fig. 3D).

Finally, if overall age effects contained a stable structure corresponding to the motor functional chain, unsupervised clustering or principal component analysis (PCA) would be expected to approximately recover the relative organization of Control, Transmission, and Peripheral tissues. Neither analysis revealed a clear hierarchical arrangement, with tissues from the three functional levels remaining intermingled (Fig. 3E, F). Together, these results indicate that transcriptional aging of the motor system does not follow a stable central-to-peripheral temporal sequence and does not form an overall continuous structure consistent with the motor functional chain.

### Age-dependent reorganization of cross-tissue transcriptional coordination

Given the absence of a stable transcriptional aging sequence along the motor functional chain, we next examined how cross-tissue coordination changed dynamically with age. Within the age range shared across all tissues, mean cross-tissue transcriptional synchrony progressively increased from around age 30, reached its highest level around age 50 (mean ρ = 0.185), and subsequently declined toward age 70. Gene-level bootstrap analysis supported the decline from the 55–65-year to the 65–75-year window (Fig. 4A). This dynamic was accompanied by a weakening of the functional-chain structure. The negative association between functional-chain distance and window-specific transcriptional synchrony was most pronounced around ages 30 and 40 (ρ = −0.414 and −0.406, respectively) and weakened thereafter (Fig. 4B), indicating a progressive attenuation of the functional-chain gradient with age.

**Fig. 4:**
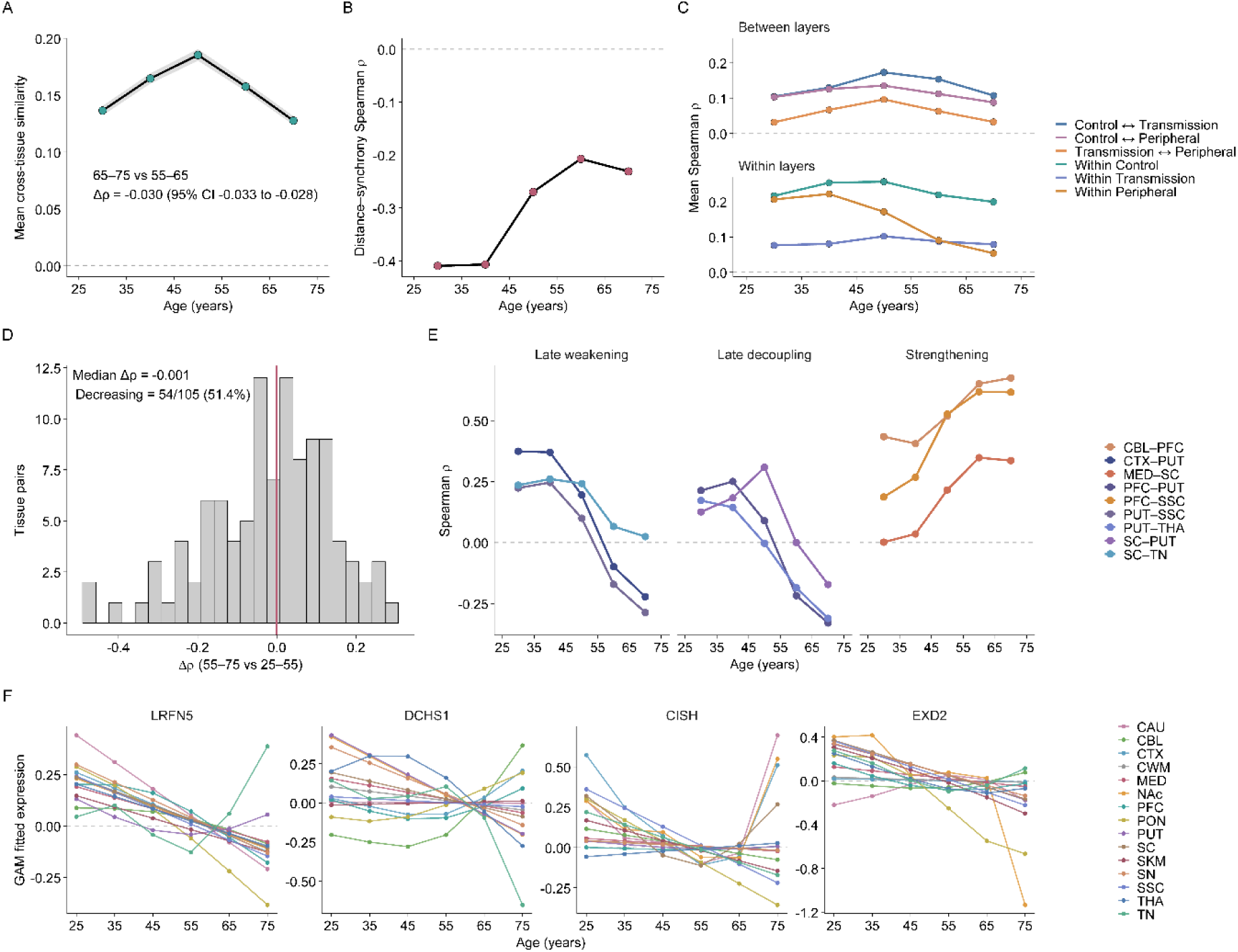
Age-dependent reorganization of cross-tissue transcriptional coordination. **A** Mean cross-tissue transcriptional synchrony across age windows. Shaded regions indicate gene-level bootstrap 95% confidence intervals. **B** Spearman correlation between functional-chain distance and cross-tissue transcriptional synchrony across age windows. **C** Age-dependent changes in mean transcriptional synchrony within and between functional levels. **D** Distribution of changes in tissue-pair synchrony in later life relative to early-to-mid adulthood; the red line indicates the median. **E** Age-dependent dynamics of representative tissue connections showing late weakening, late decoupling, or strengthening. **F** Cross-tissue age trajectories of representative genes showing transcriptional reorganization.

Similar but quantitatively distinct dynamics were observed across functional levels. Synchrony between all three pairs of functional levels increased toward midlife and subsequently declined. In particular, Transmission–Peripheral synchrony reached a relatively high level around age 50 and decreased markedly by age 70 (mean ρ = 0.096 versus 0.032). Synchrony within the Control level also declined (mean ρ = 0.257 versus 0.199), whereas changes within the Transmission level were comparatively modest. Within the Peripheral level, synchrony began to decline earlier and continued to decrease across later age windows (ρ = 0.222, 0.171, 0.090, and 0.054; Fig. 4C).

When mean synchrony in later life (55–75 years) was compared with that in early-to-mid adulthood (25–55 years), 54 of 105 tissue pairs (51.4%) showed decreased synchrony, indicating an overall tendency toward weaker cross-tissue coordination in later life. However, the magnitude and direction of these changes varied substantially among tissue pairs (Fig. 4D; Supplementary Fig. 2). The complete pairwise profiles further showed that late-life changes were distributed heterogeneously across the motor system rather than representing a uniform system-wide decline (Supplementary Fig. 2).

Among individual tissue connections, several of the most pronounced late-life decreases involved PUT, including CTX–PUT, PFC–PUT, PUT–SSC, PUT–THA, and PUT–SC. PFC–PUT and PUT–SC, in particular, shifted from positive synchrony around midlife toward negative synchrony in later life. SC–TN also weakened markedly after age 55. By contrast, PFC–SSC, MED–SC, and CBL–PFC showed progressive strengthening with age (Fig. 4E; Supplementary Fig. 2). These patterns indicate that late-life changes do not reflect uniform loss of coordination across the system, but rather a reorganization in which some connections become decoupled while others are maintained or strengthened.

Representative genes illustrated corresponding forms of cross-tissue reorganization. LRFN5 showed relatively high cross-tissue consistency earlier in adulthood followed by pronounced divergence in later life. DCHS1 showed an initial convergence of tissue trajectories followed by renewed divergence at older ages. CISH exhibited highly concordant decreases during early-to-mid adulthood but developed both directional and magnitude divergence in later life, whereas EXD2 was characterized mainly by late-life divergence in effect magnitude against a background of broadly shared decreases (Fig. 4F).

### Age-related pathway reorganization

We next examined how transcriptional reorganization was manifested at the pathway level. Across the common age range, 564∼2,355 significant GO-BP terms were identified in individual tissues. Functional enrichment was predominantly negative at earlier ages, with negatively enriched pathways accounting for 91.0% and 84.6% of significant pathways around ages 30 and 40, respectively. Positive processes became more prominent around age 50, the two directions became more balanced around age 60, and negative enrichment again predominated around age 70 (70.7%; Fig. 5A).

**Fig. 5:**
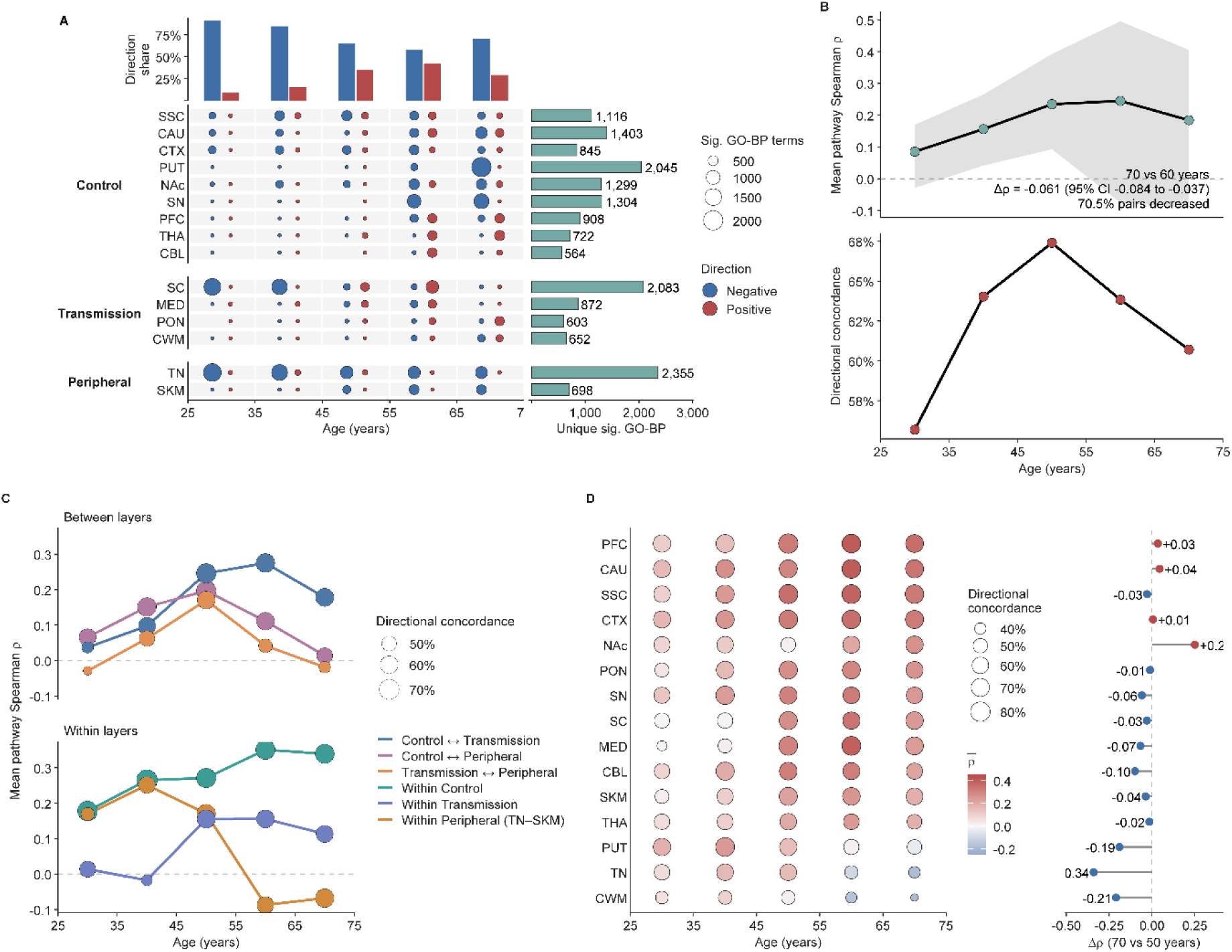
Age-related pathway reorganization and late-life cross-tissue decoupling in the motor system. **A** Direction and number of significant GO-BP terms across tissues and age windows. The stacked bars at the top show the overall proportions of positively and negatively enriched terms at each age, and the bars on the right show the number of unique significant GO-BP terms identified in each tissue across the common age range. **B** Age-dependent changes in system-wide cross-tissue pathway coordination. The upper panel shows the mean Spearman correlation of pathway NES values across tissue pairs, with the shaded region indicating the interquartile range; the lower panel shows mean directional concordance among significant pathways. **C** Changes in pathway synchrony within and between functional levels; point size indicates directional concordance. **D** Tissue-specific reorganization of pathway coordination. The panel on the right shows the change in synchrony age (Δρ).

Cross-tissue pathway coordination progressively strengthened through midlife and weakened in later life. Age-related responses of the same pathways across tissue pairs became increasingly similar toward midlife, reaching the highest level around age 60 before declining around age 70, with mean Spearman correlations of 0.085, 0.156, 0.234, 0.245, and 0.184 across successive age windows. Directional concordance showed a similar pattern but peaked earlier, around age 50, and subsequently declined, with mean concordance among significant pathways decreasing from 67.4% to 63.8% and 60.7%. Comparing the windows centered around ages 60 and 70, pathway coordination decreased in 70.5% of tissue pairs, with a mean Δρ of −0.061 (bootstrap 95% CI, −0.084 to −0.037), supporting a system-wide weakening of cross-tissue functional coordination in later life (Fig. 5B).

This late-life decoupling was concentrated primarily across functional levels and in connections involving the Peripheral level. Within-Control pathway synchrony remained relatively high at older ages, whereas Control–Peripheral and Transmission–Peripheral synchrony declined markedly, from mean correlations of 0.197 to 0.014 and 0.171 to −0.019, respectively. Within the Peripheral level also showed pronounced reorganization: mean pathway correlation decreased from 0.171 around age 50 to −0.087 around age 60 and remained low at −0.068 around age 70. Despite this loss of correlation, NES directions among pathways significant in both tissues remained largely concordant at the two later windows (84.2% and 86.0%, respectively), suggesting that the decoupling primarily reflected reorganization of pathway ranking and response magnitude rather than a global reversal of direction (Fig. 5C).

Among individual tissues, TN showed the most pronounced late-life change. Its mean pathway coordination with other tissues decreased from 0.154 around age 50 to −0.187 around age 70 (Δρ = −0.341), while directional concordance among significant pathways declined from 61.4% to 40.6% (Fig. 5D).

### Representative functions of age-related reorganization

A total of 869 robust GO-BP terms were grouped into 441 functional modules based on semantic similarity (Supplementary Fig. 3). Negative modules recurrent across multiple age windows and functional levels predominantly involved energy metabolism. These recurrent patterns indicate that mitochondrial energy production and associated metabolic processes constitute a persistent functional background of age-related reorganization across the motor system (Fig. 6A).

**Fig. 6:**
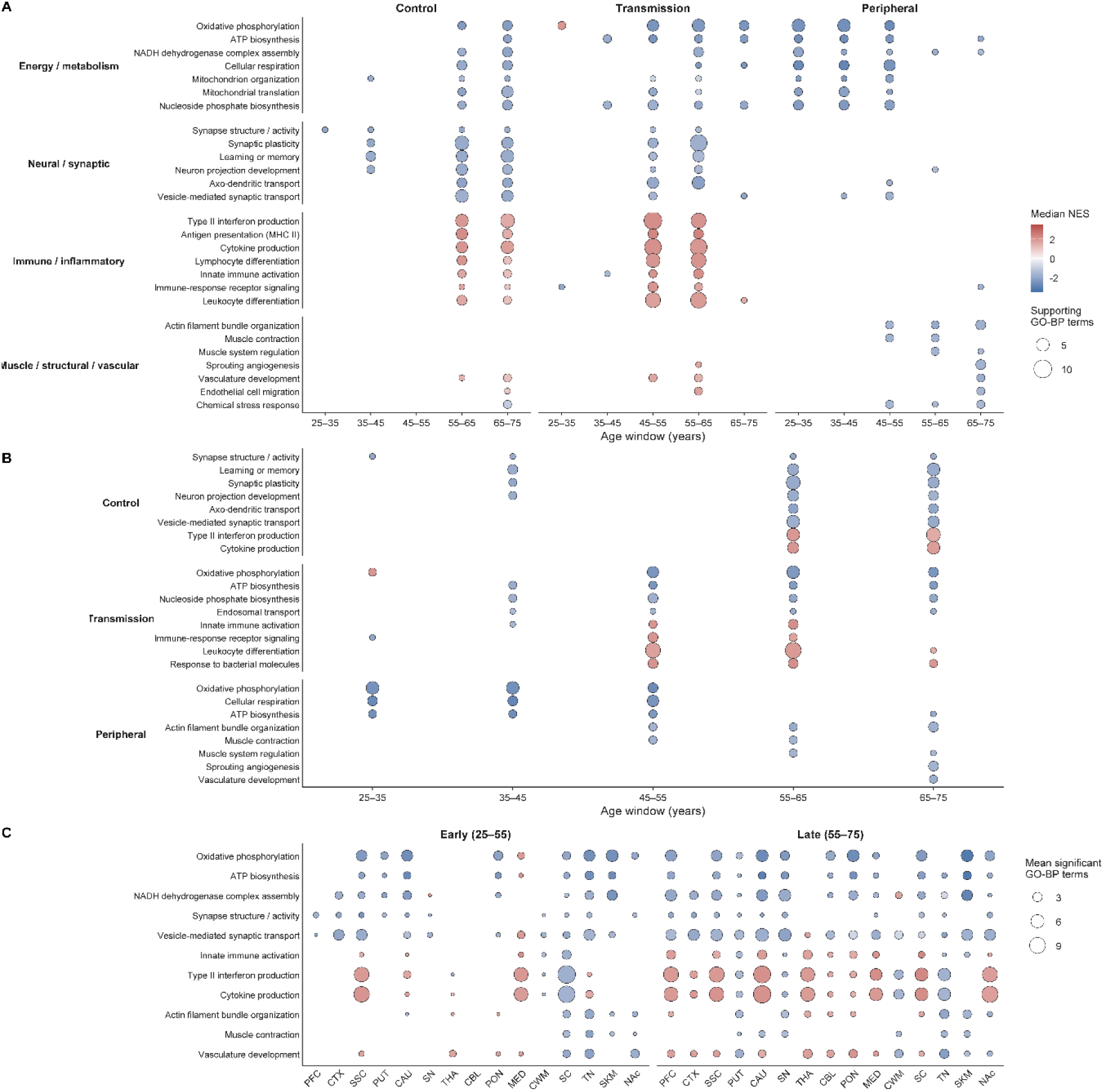
Functional-level and tissue-specific patterns of age-related pathway reorganization in the motor system. **A** Reorganization of representative functional modules across functional levels and age windows. Bubble size indicates the number of robust GO-BP terms supporting each module; blank positions indicate the absence of a coherent functional module for the corresponding functional level and age window. **B** Age-dependent dynamics of representative functional modules within the Control, Transmission, and Peripheral levels. **C** Distribution of representative functional modules across individual tissues during early (25–55 years) and late (55–75 years) periods. Bubble size indicates the mean number of significant GO-BP terms across age windows; blank positions indicate that no significant member GO-BP terms were detected for the corresponding tissue and age period.

At the Control level, functional changes were concentrated primarily in neuronal and synaptic processes. Negative age-related responses were already evident at earlier ages in synaptic and neuronal processes, including synaptic regulation, learning and memory, and neuron projection development. After age 55, negative responses persisted in synaptic plasticity, neuronal projection, and axonal and synaptic transport. In parallel, positive enrichment of immune-related processes emerged during mid-to-late life, including interferon signaling, antigen processing and presentation, cytokine production, and lymphocyte differentiation, indicating the coexistence of negative neuronal responses and increasing immune-related activity (Fig. 6A, B).

Functional reorganization at the Peripheral level showed a clear age-dependent progression. Between ages 25 and 55, negative responses predominantly involved mitochondrial energy metabolism and related metabolic processes. After age 55, the affected functions expanded to include cytoskeletal organization and muscle-related processes. By ages 65–75, additional changes emerged in vascular and stress-related processes. Thus, Peripheral reorganization progressively expanded from predominantly bioenergetic and metabolic processes at earlier ages to muscle structural, vascular, and stress-related processes in later life (Fig. 6A, B).

The Transmission level showed an intertwined pattern of metabolic, neural, and immune reorganization. Energy metabolism and intracellular transport processes recurred across multiple age windows and were predominantly negatively enriched, whereas immune-related functions became more prominent after midlife, including innate immune activation, immune signaling, and leukocyte-related processes. Several functional modules also showed age-dependent directional shifts. For example, oxidative phosphorylation changed from positive enrichment at earlier ages to negative enrichment during mid-to-late life, whereas innate immune activation and immune-response receptor signaling shifted from negative enrichment at earlier ages to positive enrichment around midlife. These patterns indicate that the Transmission level undergoes functional reorganization characterized by concurrent metabolic attenuation and immune activation during mid-to-late life (Fig. 6A, B). These representative functional modules also showed marked differences across individual tissues and between early and late age periods (Fig. 6C).

## Discussion

Molecular aging proceeds at different rates across tissues while also exhibiting shared age-related patterns [5,6,9,10]. The contribution of age to gene-expression variance can differ by more than 20-fold across tissues [5], and different organs within the same individual can exhibit distinct biological ages [9]. At the same time, altered intercellular communication has been recognized as a core hallmark of aging [8], suggesting that changes in relationships between tissues may themselves constitute part of the aging process. This issue is particularly relevant to the motor system because movement is not the output of a single tissue but depends on distributed circuits involving the cortex, basal ganglia, brainstem, spinal cord, peripheral nerves, and skeletal muscle [1,2]. We therefore analyzed age-related changes across 15 tissues within a unified motor functional chain to systematically examine tissue specificity, temporal ordering, and cross-tissue coordination of transcriptional aging.

Our results first clarify how shared and heterogeneous features of aging can coexist. Across the motor system, 87.1% of ARGs were significant in at least two tissues, yet sharing the same ARGs did not imply that different tissues aged in the same manner (Fig. 2). Even when large numbers of ARGs were shared, the direction, magnitude, and overall correlation of age effects could differ substantially across tissues. This is consistent with previous evidence for tissue-and cell-type-specific aging responses [5,6,26–28]. Human brain and skeletal muscle atlases have shown that aging affects a broad range of common biological processes, while the magnitude of these responses differs markedly across cell populations [26–28]. For example, age-related reductions in synaptic gene expression in the brain vary across cell types [26], whereas aged skeletal muscle exhibits distinct responses across muscle fibers, muscle stem cells, and neuromuscular junction-associated cell populations [27,28]. Our findings therefore suggest that tissue-specific aging within the motor system does not necessarily arise primarily from entirely distinct sets of aging-related genes, but may instead reflect differences in the weighting, direction, and temporal organization of shared age-related programs across tissues. However, these tissue-specific responses did not form a stable central-to-peripheral aging sequence. The timing of major transcriptional changes was nearly indistinguishable across the Control, Transmission, and Peripheral levels, and functional distance did not explain similarity in age effects between tissues (Fig. 3). Although the motor system has a clear functional direction from central command to peripheral execution, motor control is implemented through extensive parallel, feedback, and recurrent circuits, and interactions among the basal ganglia, brainstem, cerebellum, and cortex are not organized as a simple serial pathway [1,2]. Multi-organ studies have likewise demonstrated substantial asynchrony in aging across organs and complex inter-organ association networks rather than a single unified systemic aging timetable [6,9,10]. Thus, the direction of motor information flow cannot be directly translated into a direction of molecular aging propagation. This may reflect fundamental differences between physiological function and transcriptional organization, as well as distinct temporal properties of aging itself.

More important than a fixed sequence was the observation that cross-tissue relationships themselves changed with age. Gene-level cross-tissue synchrony progressively increased during early-to-mid adulthood, reached relatively high levels around midlife, and subsequently declined in later life. This decline was not uniform across the system: some tissue connections weakened, whereas others were maintained or even strengthened (Fig. 4). This is consistent with an emerging view that aging alters not only the expression of individual genes but also the coordination among molecular processes [7,11–13]. Leote et al. reported that aging weakens gene–gene relationships among some fundamental cellular processes while strengthening others [11]. Our results extend this concept to a higher organizational level: rather than examining relationships among genes within a single tissue, we show that age-related responses across different motor-system tissues progressively lose their correspondence. In this context, late-life decoupling is better understood as a reconfiguration of pre-existing cross-tissue relationships than as a uniform collapse of coordination.

This age-dependent pattern may reflect a changing balance between shared systemic pressures and tissue-specific local adaptation. With advancing age, inflammatory, metabolic, endocrine, and circulatory factors can act simultaneously on multiple tissues [8,29]. Physical activity itself also establishes bidirectional communication between muscle, brain, and other organs through myokines, metabolites, and other exerkines [30,31]. The increase in cross-tissue coordination during mid-adulthood may therefore partly reflect convergent responses to shared age-related pressures or adaptive signals. In later life, however, increasing divergence in metabolic reserve, cellular composition, repair capacity, and local microenvironment across tissues may make such coordinated responses progressively more difficult to maintain. Importantly, the coordination measured here represents statistical correspondence in age effects rather than direct evidence of inter-tissue signaling.

The locations of decoupling also showed features specific to the motor system. At the gene level, many of the strongest late-life declines in synchrony involved PUT. Normal aging is known to affect striatal dopamine signaling [32], frontostriatal structural and functional connectivity, and sensorimotor prediction and motor control [4]. The decline in PUT-related connections may therefore reflect changes in the integrative role of the basal ganglia within the aging motor network rather than simply local transcriptional deterioration in PUT. Conversely, connections such as PFC–SSC, MED–SC, and CBL–PFC were maintained or strengthened with age. Functional imaging studies similarly suggest that normal motor aging involves both reduced efficiency in some networks and task-dependent recruitment of additional regions and network reorganization [33]. These strengthened connections may therefore represent compensatory or reorganized coordination required to maintain function. Peripheral findings revealed another form of decoupling. TN showed the largest late-life decline in pathway coordination, and TN–SKM synchrony shifted from positive to negative values; however, more than 80% of pathways significant in both tissues retained concordant NES directions (Fig. 5). This suggests that peripheral nerve and muscle do not simply age in opposite directions, but may continue to respond to similar age-related pressures while progressively diverging in response magnitude and functional priority. Two recent human skeletal muscle aging atlases have reported reduced innervation, neuromuscular junction remodeling, changes in terminal Schwann cells, and repeated denervation–reinnervation with aging, accompanied by muscle-fiber remodeling and regenerative responses [27,28]. Increased abundance of NMJ-associated nuclei has also been proposed to contribute to reinnervation [28]. These observations are consistent with the TN–SKM pattern of preserved directional concordance but declining coordination: peripheral nerve and muscle may continue to activate related maintenance or stress-response programs while becoming increasingly mismatched in compensatory capacity and response intensity. Given that loss of muscle strength and impaired innervation are both major components of sarcopenia [3,27,28], altered nerve–muscle molecular matching may be more closely related to motor functional decline than changes in muscle alone.

Pathway-level analyses further indicated that late-life decoupling was accompanied by increasing divergence in aging states across functional levels (Fig. 6). The Control level was characterized primarily by persistent negative responses in neuronal, axonal, and synaptic processes together with increased immune-related activity during mid-to-late life. The Peripheral level showed earlier declines in mitochondrial and energy-metabolic processes, followed by broader remodeling involving muscle structure, the cytoskeleton, and vascular functions. The Transmission level displayed more pronounced stage-specific dynamics, with metabolic processes generally shifting toward negative responses while selected immune processes increased after midlife and some pathways exhibited age-dependent directional switching. Thus, late-life system-level decoupling does not appear to reflect simultaneous entry of all tissues into a single aging state, but rather the progressive emergence of distinct functional remodeling programs across different levels of the motor system. This further suggests that motor aging cannot be fully explained by changes in either the brain or skeletal muscle alone.

This study has several limitations. Although GTEx includes multiple tissues from many of the same donors [14], GEO datasets were largely derived from different studies and individuals. Our age trajectories therefore reflect cross-sectional population-level patterns rather than longitudinal aging within individuals and cannot directly capture within-individual physiological coupling across tissues. To harmonize GTEx and GEO and ensure comparability across age windows, we used 10-year age intervals; consequently, our analyses are better suited to identifying stage-dependent changes than precise transition ages, which require validation using exact ages and longitudinal data. In addition, bulk-tissue age effects reflect both shifts in cellular composition and cell-intrinsic transcriptional changes [26–28], which were not resolved at single-cell resolution.

Overall, our findings support a systems-level view of aging across motor-related tissues. Tissues throughout the motor system share broad age-related molecular changes, but these changes do not progress sequentially along the central-to-peripheral functional chain. Instead, responses to shared age-related changes become increasingly differentiated across tissues with advancing age, culminating in selective late-life decoupling of transcriptional and functional coordination across the motor system.

## Acknowledgements

H.N. acknowledges support from the Japan Science and Technology Agency (JST) through the Support for Pioneering Research Initiated by the Next Generation (SPRING), under the Program for Development of Next-Generation Front-Runners with Comprehensive Knowledge and Humanity at the Institute of Science Tokyo.

