## Supplementary Figures for "Age-Related Remodeling of Cross-Tissue Transcriptional Coordination in the Human Motor System"

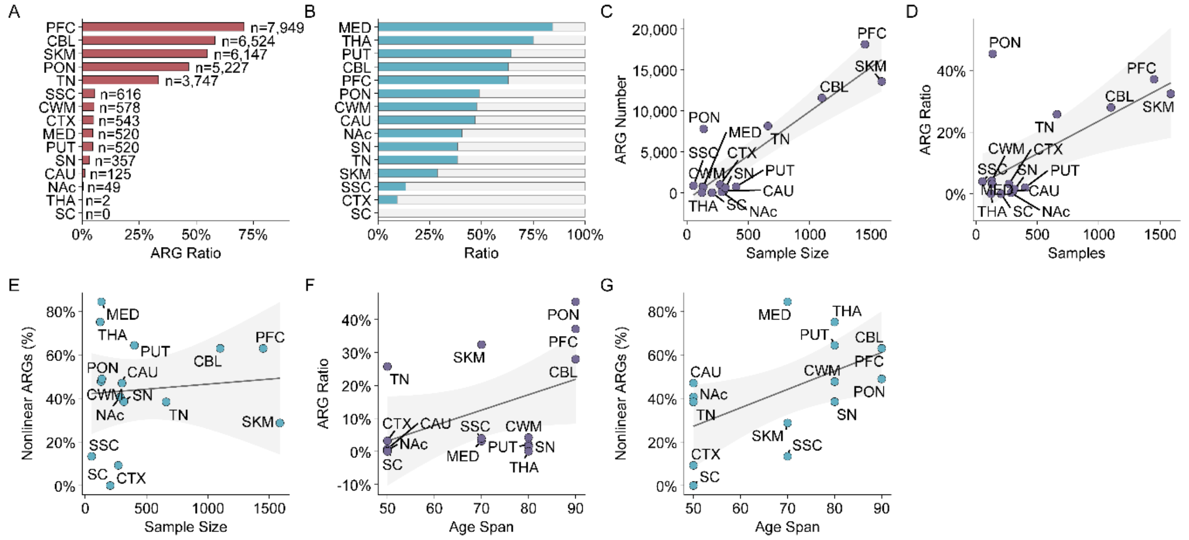

**Supplementary Fig. 1: Robustness and technical assessment of age-related transcriptional patterns.**

**A** ARG proportion based on the common-gene background. **B** Proportions of linear and nonlinear ARGs. **C–E** Relationships of sample size with ARG number, ARG proportion, and nonlinear ARG proportion, respectively. **F, G** Relationships of age span with ARG proportion and nonlinear ARG proportion, respectively. Lines indicate linear regression fits and shaded regions indicate 95% confidence intervals.

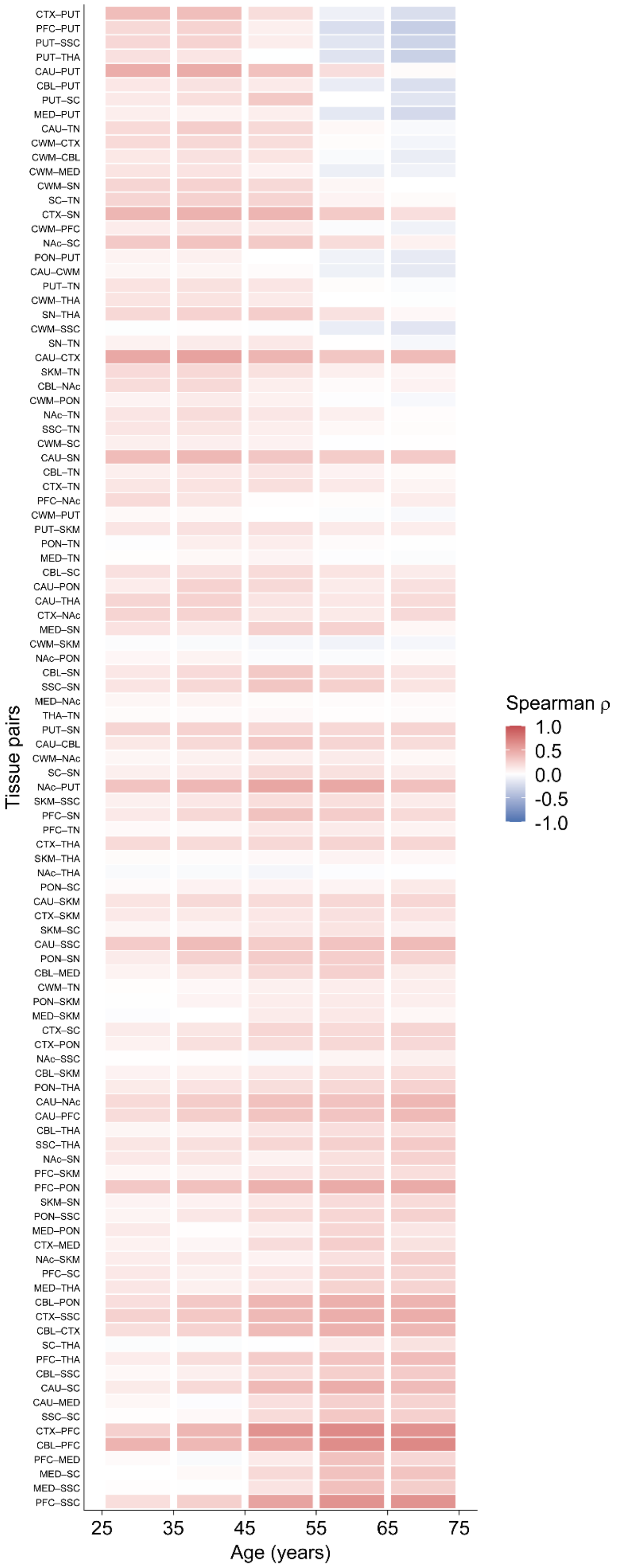


**Supplementary Fig. 2: Age-dependent transcriptional synchrony across all tissue pairs.** Heatmap of Spearman correlations (ρ) of transcriptional changes across 105 tissue pairs and five consecutive age windows. Tissue pairs are ordered by the change in mean synchrony between later life (55–75 years) and early-to-mid adulthood (25–55 years).

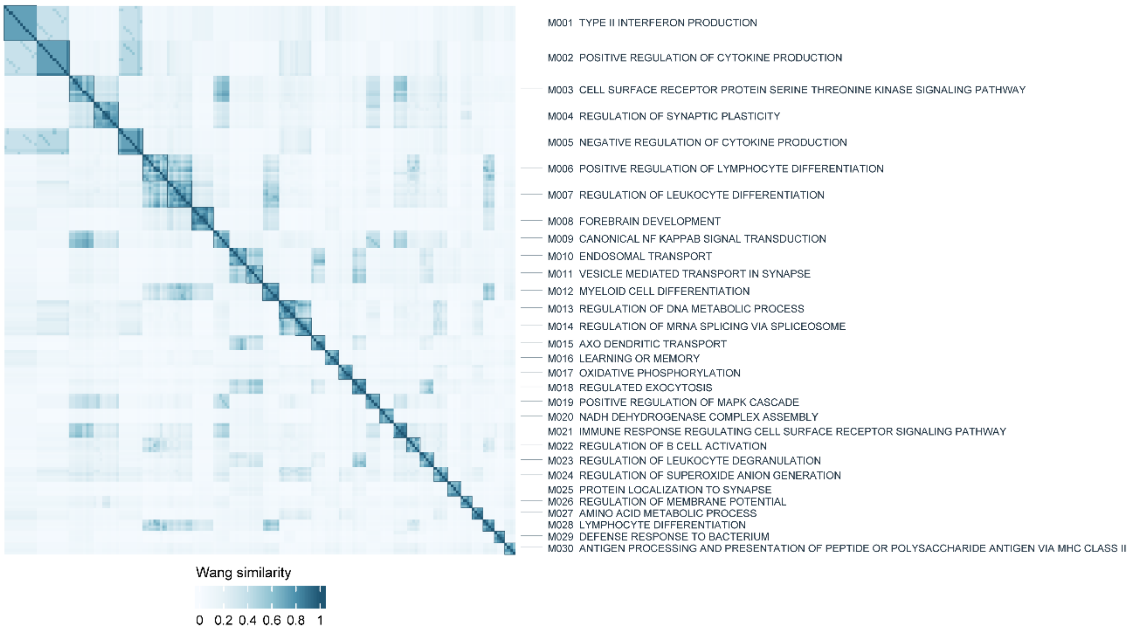


**Supplementary Fig. 3: Semantic similarity of robust GO-BP terms.**
Heatmap of pairwise Wang semantic similarity among top 30 robust GO-BP terms. Terms are grouped into semantic modules using a similarity threshold > 0.7 and edge-betweenness community detection; M001–M030 indicate the largest modules and their representative terms.
